# Fine-Scale Structural Information Substantially Improves mRNA Therapeutic Stability Prediction

**DOI:** 10.1101/2025.08.15.670605

**Authors:** Soon Yi, Sara E. Ali, Yashrajsinh Jadeja, J. Wade Davis, Mihir Metkar

## Abstract

The success of COVID-19 mRNA vaccines has made the in-solution stability optimization of mRNAs a key objective. However, we still lack a complete understanding of sequence metrics that influence mRNA in-solution stability. RNA secondary structure plays a critical role in protecting against hydrolysis, the primary degradation pathway under storage conditions. Yet, the structural metrics that best guide stability-focused mRNA design remain incompletely defined. Global metrics like minimum free energy and average unpaired probability have improved mRNA stability but fail to capture local structural variation relevant to RNA degradation. We demonstrate that base-pairing log odds provide fine-scale, orthogonal insight that complements global metrics and improves stability modeling. Further, by combining local and global features into a parsimonious four-feature regression model, dubbed **St**ability **R**egression **A**nalysis using **N**ucleotide-**D**erived features (STRAND), we achieve a greater than two-fold reduction in prediction error compared to existing machine learning and deep learning approaches and demonstrate robust generalization across diverse transcript contexts. This compact and interpretable model provides an accurate and reliable framework for predicting mRNAs in-solution stability.

## INTRODUCTION

Optimizing therapeutic mRNA requires not only effective protein expression but also in-vial stability that resists degradation during storage and delivery. Although the rapid mRNA-based vaccine development has highlighted the platform’s potential^1^, mRNA instability remains a critical barrier to broader deployment, particularly in settings lacking reliable cold-chain infrastructure such as developing countries^2^. Degradation primarily occurs through hydrolysis and transesterification^3,4^ reactions in-solution during storage in-vial, typically initiating at structurally accessible nucleotides within unpaired loops or flexible regions^5,6^. Consequently, solution conditions such as temperature, pH, and ionic strength can drastically reduce mRNA half-life, thereby limiting therapeutic efficacy and significantly increasing manufacturing complexity and costs^7^.

A key mRNA degradation determinant is its secondary structure, which arises from intramolecular base pairing. Base-paired regions can shield vulnerable sites from chemical or enzymatic attacks, extending RNA stability^6^. Accordingly, rational mRNA design has focused on engineering sequences that adopt favorable structural conformations. However, this design process requires reliable metrics that are predictive of RNA stability^8,9^.

Traditional metrics such as free energy (FE), average unpaired probability (AUP), and GC content have been widely used to guide mRNA optimization^10,11^. Free energy minimization identifies the most thermodynamically stable structure an RNA can adopt^12,13^; AUP captures the average likelihood that nucleotides remain unpaired; and GC content correlates with duplex stability due to stronger G-C base pairing^8,14^. These metrics are computationally efficient and biologically relevant, and they have been integrated into mRNA design tools^15,16^ such as CDSFold^17^, RiboTree^18^, LinearDesign^19^ and DERNA^20^. However, despite their usefulness, these metrics do not fully explain observed degradation differences between mRNA sequence variants as reflected by their modest Spearman correlation coefficient (<0.6)^9^, suggesting they fail to capture important aspects of RNA structural ensembles^9^.

Large-scale efforts have attempted to model mRNA degradation using both statistical and deep learning approaches. For example, the dual crowd-sourcing study by Wayment-Steele et al.^21^ combined human- and machine-designed sequences to benchmark predictive models of degradation, including those incorporating experimental selective 2’-hydroxyl acylation analyzed by primer extension (SHAPE) data^6,22,23^. Despite access to thousands of synthetic constructs and high-resolution structural probing data, models achieved only moderate predictive accuracy when applied to full-length mRNAs, with Spearman correlation coefficients (rho) ranging from 0.36 to 0.48^21^. These results reflect the inherent difficulty of modeling mRNA degradation, which likely depends on a combination of global and local structural features not fully captured by existing metrics.

To address this limitation, we propose a new metric: the base pairing log-odds (LO), which quantifies the base-pairing likelihood at each position across the structural ensemble by comparing the odds of being paired to unpaired. Unlike base-pairing probability^24,25^, which is bounded between 0 and 1 and compresses extreme values, the log-odds transformation expands the dynamic range^26^, particularly for high and low probabilities, thereby amplifying meaningful structural differences that might otherwise be obscured. LO reflects the local base-pairing probability distribution, offering finer resolution than global metrics like FE or AUP. By analyzing mRNAs with similar global scores but divergent in-solution half-lives, we show that LO captures functional differences not apparent in traditional metrics. Finally, we demonstrate that incorporating LO into a compact four-feature regression model, <u>St</u>ability <u>R</u>egression <u>A</u>nalysis using

Nucleotide-Derived features (STRAND), significantly improves mRNA stability prediction over existing approaches and generalizes across diverse transcript contexts, providing an interpretable framework for stability-focused mRNA design.

## RESULTS

### Traditional RNA Structural Metrics Partially Explain In-Solution Stability

A major goal in RNA therapeutic design is to extend mRNA half-life in-vial^27^. Existing studies have attempted to improve stability using sequence-derived structural metrics, such as FE and AUP, which provide approximations of RNA secondary structure^18^. However, whether these metrics alone are sufficient to predict mRNA degradation rates remains an open question^9^.

To assess the relationship between RNA structure and in-solution degradation, we examined a dataset of Nano Luciferase (NLuc), enhanced green fluorescence protein (eGFP), and multi-epitope vaccine (MEV) mRNAs with measured half-lives in-solution (**Figure 1A**)^9^. NLuc sequences were selected for further analysis due to their broad half-life range and the largest number of available sequence variants (N = 69), making them an ideal test set for evaluating structural predictors.

**Figure 1.**
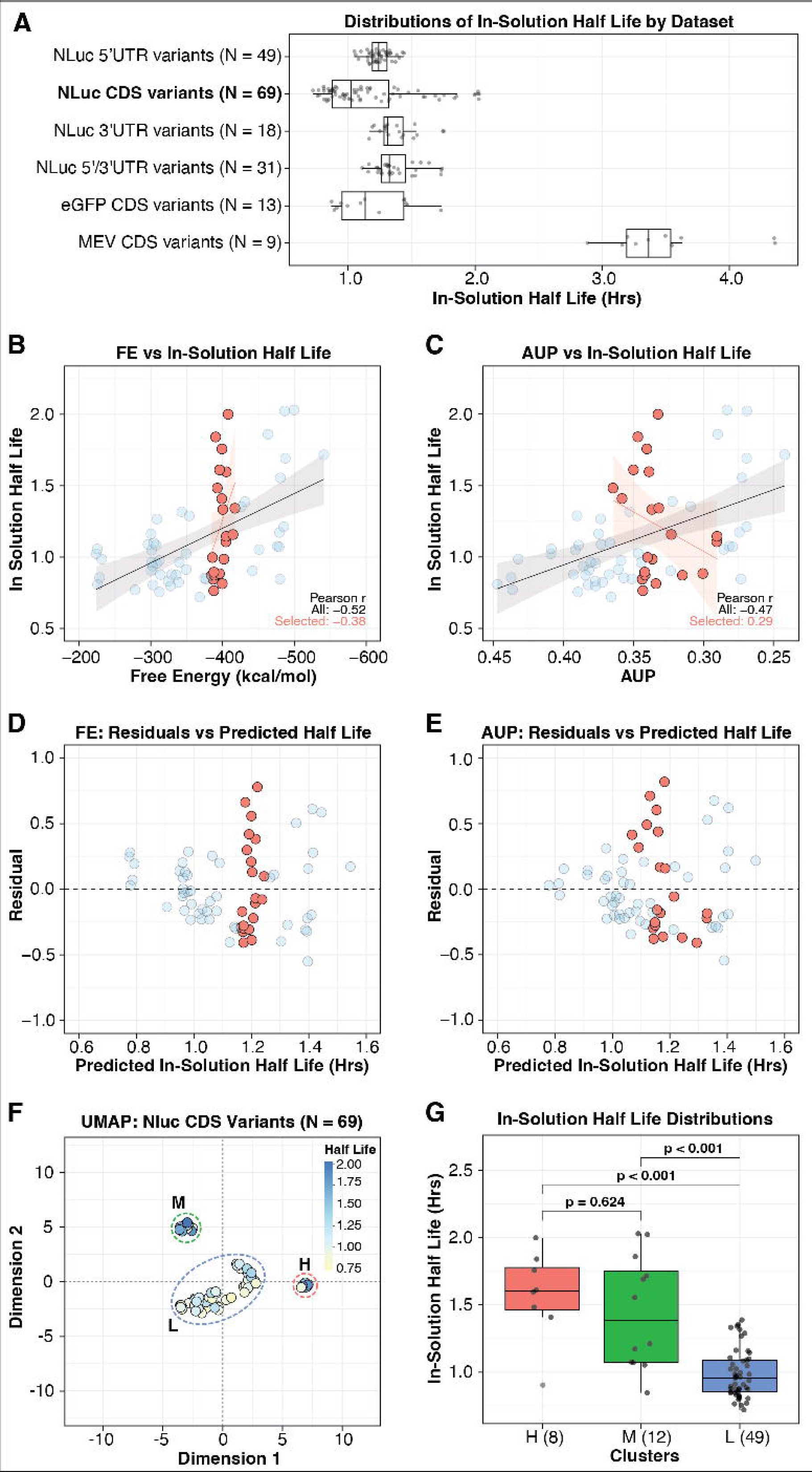
In-solution half-life distributions across protein mRNA variants and structure–function analysis in the Nano Luciferase set. (A) A boxplot distribution of in-solution half-lives for mRNA variants of different proteins. The box represents the interquartile range, the horizontal line indicates the median, and whiskers denote 1.5X the interquartile range; individual sequences are plotted as points. The NanoLuc (NLuc; bold) CDS variants were selected for further analysis due to their wide range of half-lives and large number of sequences. (B) Correlation of FE with half-life in the NLuc dataset. Red-highlighted sequences show similar FE but divergent half-lives. Pearson r for the entire dataset (black) and for sequences with similar FE (red) are shown. (C) Same as (B) but for AUP. (D) Residual plots for FE showing increased error for sequences with higher half-lives. (E) Same as (D) but for AUP. (F) UMAP projection of NLuc sequences based on global structural features (FE, AUP, and GC content), revealing three clusters. Clusters are labeled as high (H), medium (M), and low (L) based on their in-solution half-lives. (G) Boxplot distribution of half-lives for sequences in each UMAP-defined cluster in (F), with pairwise statistical comparisons shown. The Mann-Whitney test was used to calculate p-values. The number of sequences within each cluster is noted in parentheses.

Across the NLuc dataset, we observed moderate negative correlations between half-life and FE (Pearson r = −0.52) as well as AUP (Pearson r = −0.47) (red and blue points in **Figure 1B, 1C**). These correlations suggest that mRNAs with lower FE and AUP, indicative of more stable secondary structures, tend to degrade more slowly. However, while these metrics broadly capture structural trends, they fail to fully account for the variability in observed degradation rates. For example, the NLuc sequence variants with FE around −400 kcal/mol (N = 21, NLuc-21 set) showed a lower correlation than the global comparison, with Pearson correlation values of −0.38 and 0.29, respectively (**Figure 1B, 1C**, highlighted in red). Moreover, linear regression models using FE and AUP as predictors exhibited high residual errors, particularly at higher values of either metric (**Figure 1D, 1E**). We also noted similar observations for eGFP CDS variants (N = 13; **Figure S1**), in which eGFP sequences with FE near –500 kcal/mol show approximately 1.7-fold differences in their measured half-lives and sequences with AUP near 0.40 show approximately 2-fold differences in their half-lives. These findings suggest that the observed discrepancy between the traditional metrics and degradation rates is a general phenomenon across different coding sequence sets. While global structure formation plays a role in stability, additional sequence-dependent factors influence degradation in ways that traditional metrics fail to capture.

To further investigate this variability, we applied Uniform Manifold Approximation and Projection (UMAP) to reduce the three-dimensional structural feature space into a lower-dimensional representation.^28^ The resulting UMAP embedding, based on traditional metrics, including minimum FE normalized by transcript length (FE/length), average unpaired probability (AUP), and GC content, separated the NLuc sequences into distinct clusters (**Figure 1F**). Normalizing FE by transcript length enabled comparison across sequences of varying lengths, ensuring that the energy values reflect general structural stability rather than absolute sequence length. The resulting UMAP clusters were identified as high (H), medium (M), and low (L) based on the average in-solution half-life. Despite the apparent separation in UMAP space, clusters H and M exhibited no statistically significant difference in their half-life distributions (**Figure 1G**), underscoring the inability of traditional metrics to resolve functionally distinct stability groups. This suggests that additional sequence-derived features may be necessary to explain mRNA degradation more accurately. To explore this further, we selected two NLuc sequences with nearly identical FE and AUP values, but extreme differences in half-life, and aimed to determine whether additional features could explain their stability discrepancies.

### Local Structural Features Differentiate Sequences with Similar Global Metrics

The two sequences, X320004D1 and X930007D1, had FEs of −387.71 and −407.65 kcal/mol, a ∼5% difference, while their AUP values differed by less than 3% (0.34 vs. 0.33). Notably, X930007D1 had a higher GC content (60.03%) than X320004D1 (52.04%), though the direct effect of GC content on stability in this context remains uncertain. Their measured half-lives varied by nearly three-fold (0.72 vs. 1.99 hours, respectively) (**Figure 2A**).

**Figure 2.**
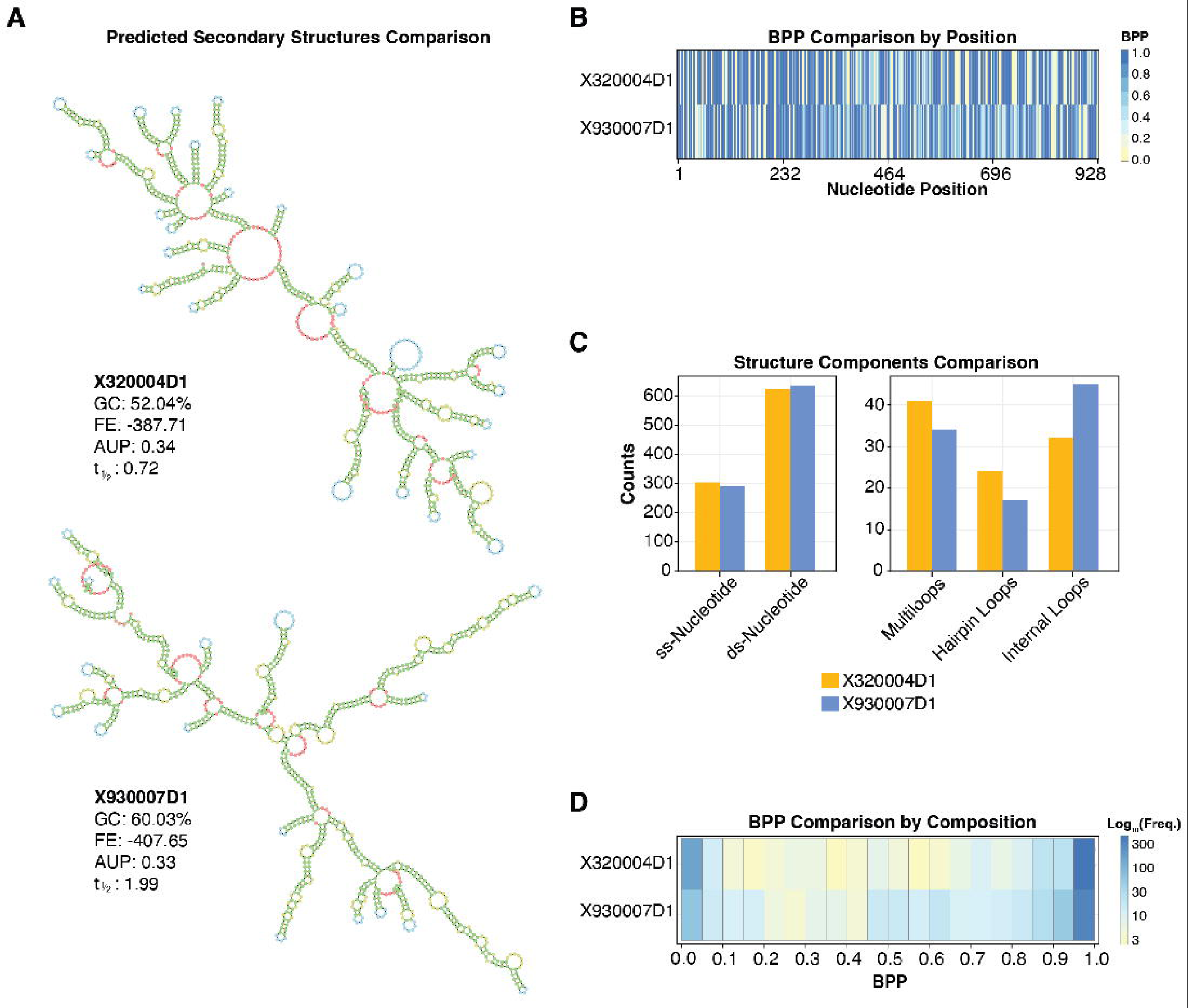
Structure comparison between two RNA sequences with similar global metrics but substantial differences in in-solution half-life. **(A)** Predicted secondary structures of two NLuc sequences, X320004D1 and X930007D1, which exhibit similar FE and AUP but differ substantially in in-solution half-life. Global structure metrics and half-life values are shown. **(B)** Heatmap of local BPP values by position for the two selected sequences. **(C)** Barplot comparing structural components between the predicted maximum expected accuracy structures, including counts of unpaired nucleotides, hairpin loops, interior loops, and multiloops. **(D)** Heatmap comparison of BPPs binned at 0.05 resolution for the two selected sequences. Color scale indicates log₁₀ frequency of positions in each bin.

To investigate the structural basis for this difference, we predicted secondary structures of both mRNAs using LinearPartition (**Figure 2A**)^29^. A heatmap visualization of per-position base-pairing probabilities (BPPs) (**Figure 2B**) illustrated that, despite similar global metrics, these sequences exhibited distinct base-pairing patterns along the sequence and BPP compositions that may contribute to their degradation differences. Specifically, X320004D1 formed a more branched secondary structure, with a higher number of multiloops (41) and hairpin loops (24), which could introduce greater structural flexibility and accessibility. In contrast, X930007D1 contained fewer multiloops (34) and hairpin loops (17) but a higher number of inner loops (45 vs. 32 in X320004D1), forming more extended helical regions (**Figure 2C**). To quantify these differences, we compared the BPP distributions of the two sequences (**Figure 2D**). The BPP distribution of X320004D1 showed a sharp bimodal pattern, with clear peaks near 0.0 (unpaired) and 1.0 (paired), suggesting more well-defined structural elements. In contrast, X930007D1 exhibited a broader, more continuous distribution, with a dominant peak near 1.0 but more intermediate BPP values, indicating a less rigid structure.

These divergences between the global metrics and stability were not unique to this sequence pair. Among sequences with higher stability (FE near −475 kcal/mol), we observed similar disconnect between global metrics and measured half-lives. For example, X310055T3 and X310005T1 showed highly distinct predicted structures (**Figure S2A**) and measured half-lives (1.08 vs 2.03 h), despite having nearly identical FE (–476.09 vs –499.53 kcal/mol), GC content (53.56 vs 52.16%), and AUP of 0.28. In addition, the two sequences showed varying structural compositions (**Figure S2B**) and BPP distributions (**Figure S2C, D**). Together, these examples demonstrate that global metrics alone do not capture the local structural variation that may underlie differences in RNA stability, motivating a more systematic analysis of BPPs across a broader sequence set.

### Base-Pairing Probability Alone Does Not Fully Explain mRNA Stability

To determine whether the structural patterns observed in X320004D1 and X930007D1 extend to a broader dataset, we analyzed 21 NLuc variants (NLuc-21; red points in **Figure 1B, C**) with similar FE and AUP values but differing in-solution half-lives. Despite having nearly identical global structural metrics, these sequences displayed distinct patterns of local base-pairing interactions (**Figure 3A, 3B**).

**Figure 3.**
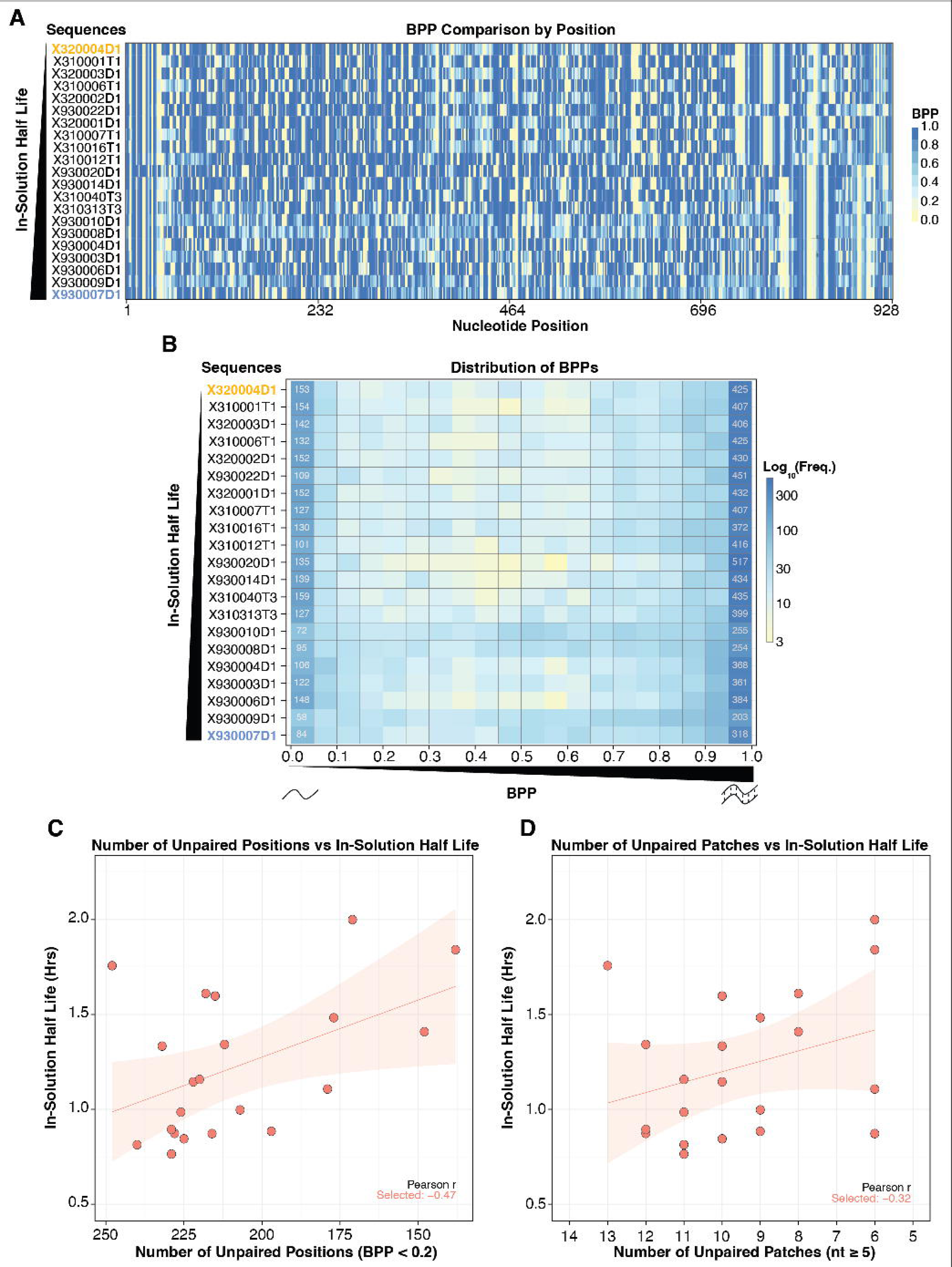
Local RNA base-pairing patterns differ among similarly structured sequences and associate with in-solution half-life. **(A)** Heatmap of local BPP values by position for sequences with similar global metrics (FE and AUP) but differing in-solution half-lives. Sequences are ordered by increasing half-life. **(B)** Heatmap comparisons of BPPs binned at 0.05 resolution across the same sequences as in **(A)**. Color scale indicates log₁₀ frequency of positions in each bin. Numbers flanking each row indicate the count of positions in the lowest and highest BPP bins. **(C)** Correlation between the number of unpaired positions (defined as BPP < 0.2) and in-solution half-life. Pearson r for the NLuc-21 sequences (red) is shown. **(D)** Correlation between the number of unpaired patches (≥5 consecutive unpaired positions with BPP < 0.2) and in-solution half-life. Pearson r for the NLuc-21 sequences (red) is shown.

A heatmap of BPPs across these NLuc-21 sequences (**Figure 3A, 3B**) revealed that, in general, each sequence possesses unique patterns of paired and unpaired regions. Some sequences exhibited more extended single-stranded regions (X320004D1), while others contained higher proportions of stable paired regions (X310012T1). Interestingly, while some sequences with higher stability displayed consistent base-pairing throughout the sequence (X930004D1), others with similar stability had more localized variations in paired and unpaired regions (X930009D1 vs X930007D1).

To quantify these differences, we examined whether the number of unpaired positions (BPP < 0.2) or the presence of long unpaired patches (≥5 nucleotides) correlated with half-life. Pearson correlation analysis showed that neither total unpaired positions (r = −0.47, **Figure 3C**) nor long unpaired patches (r = −0.32, **Figure 3D**) provided a significantly stronger correlation than FE or AUP. Moreover, our analysis showed no significant improvement in Pearson correlation with different lengths of unpaired patches (≥3 or ≥10 nucleotides; **Figure S3**).

These findings indicate that BPP alone does not account for mRNA stability differences among sequences with similar global metrics. Although BPP provides a continuous measure of pairing likelihood at each position, its bounded range (0 to 1) compresses differences at the extremes, making it difficult to resolve functionally meaningful distinctions, especially among positions with high or low pairing confidence.

Additionally, when summarized (e.g., by averaging in AUP), BPP can obscure the contribution of unpaired positions, as zero-probability sites are effectively treated as absence of signal rather than informative features. This suggests that a metric with greater dynamic range may be needed to explain the stability variation that global metrics and BPP miss.

### Log-Odds of Base-Pairing Captures RNA Stability Differences Beyond BPP

To address this limitation, we employed a base-pairing log-odds (LO) metric, which transforms BPP values into an unbounded scale (-∞ to +∞). Highly paired positions (BPP > 0.5) yield positive LO values, while strongly unpaired positions (BPP < 0.5) yield negative values. This log transformation expands the dynamic range at both extremes of the BPP distribution, amplifying differences among positions with very high or very low pairing confidence that are otherwise compressed on the 0-to-1 BPP scale.

To test whether LO captures RNA degradation patterns better than BPP, we first examined per-position LO values for X320004D1 and X930007D1 (shown in **Figure 2**). While both sequences had similar global structural metrics, LO values were distributed differently, particularly in weakly structured regions (**Figure 4A)**. Notably, X320004D1 contained a greater number of strongly negative LO positions, indicating unpaired regions that are likely more susceptible to degradation. In contrast, X930007D1 exhibited fewer such sites, suggesting fewer vulnerable positions, consistent with its longer half-life.

**Figure 4.**
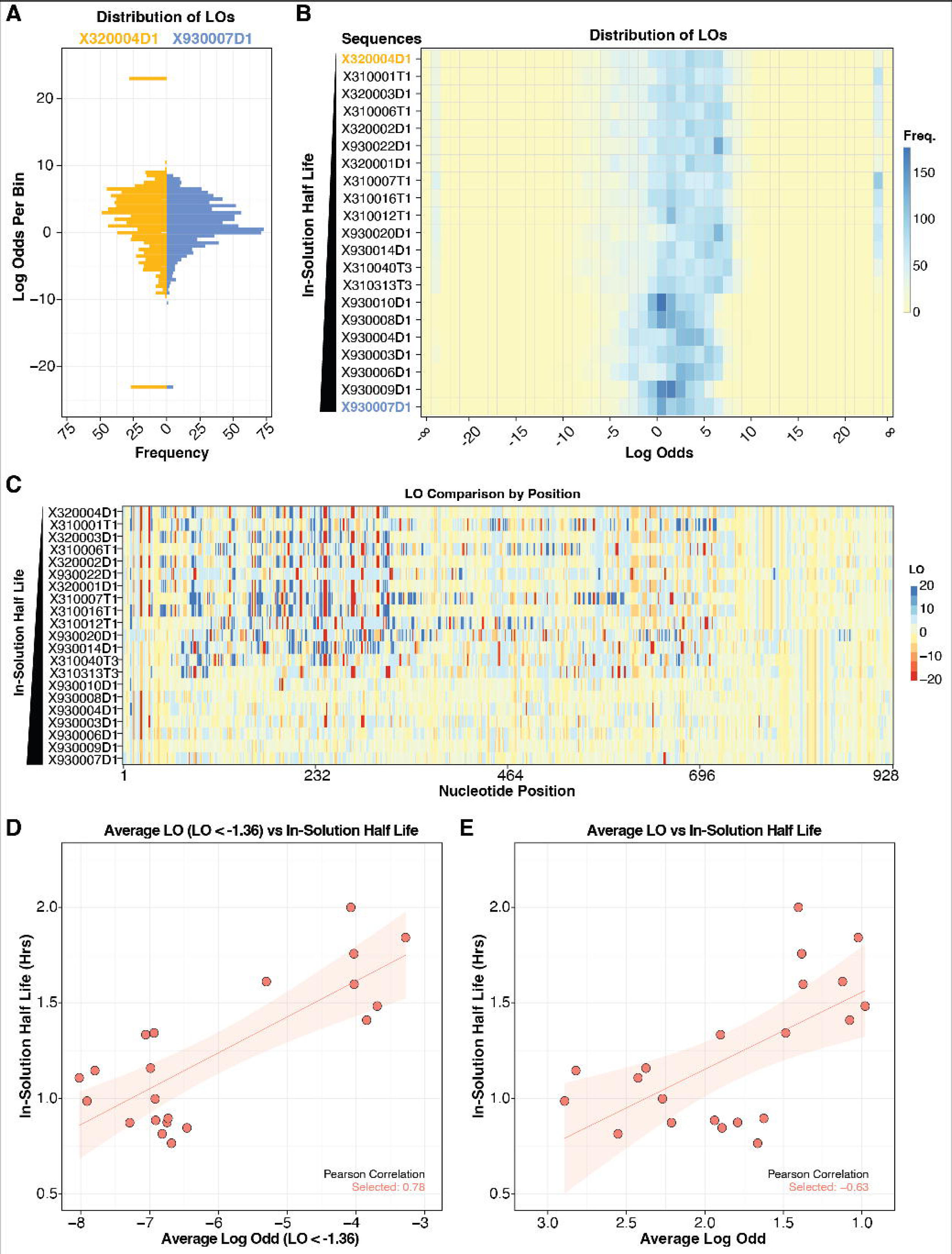
Base-pairing probability Log-odds (LO) capture local structure features predictive of RNA stability. **(A)** Histogram of LO values (binwidth = 0.5) for the two sequences (X320004D1 and X930007D1) with similar global structure metrics but divergent half-lives. **(B)** Heatmap comparisons of binned LO values across the subset of NLuc sequences (Figure 3), sorted by increasing in-solution half-life. Color scale indicates the frequency of positions in each bin. **(C)** Heatmap comparisons of LO values by position for the same sequences as in **(B)** with color scale indicating LO values and highlighting localized structural differences along the transcript. **(D)** Correlation of average LO for weakly paired regions (LO < −1.36, equivalent to BPP < 0.2) and in-solution half-life. Pearson r for the NLuc-21 sequences is shown. **(E)** Same as **(D)** but for average LO across all positions.

A heatmap visualization of LO across NLuc-21 sequences (**Figure 4B, C**) demonstrated that LO more effectively differentiated stable and unstable sequences compared to BPP. Sequences with shorter half-lives exhibited more widespread strongly negative LO values (red regions in **Figure 4C**), consistent with a greater number of confidently unpaired positions. To quantify this, we focused on weakly paired regions (LO < −1.363, derived from BPP curvature analysis) and found that the average LO (ALO) of these positions showed a strong positive correlation with in-solution half-life (Pearson r = 0.78, **Figure 4D**), indicating that sequences with less extreme negative LO values in their unpaired regions tend to be more stable.

However, when we computed the ALO across all positions, the correlation with half-life was negative (Pearson r = −0.63, **Figure 4E**). Although this opposing direction appeared counterintuitive, closer analysis of the heatmap in **Figure 4C** revealed that sequences with shorter half-lives also contained more positions with extremely positive LO values (blue regions; LO > 20). These sequences thus exhibit a more polarized LO distribution, with greater representation at both extremes, while longer-lived sequences show more moderate LO profiles overall. When averaged across all positions, the extreme positive values in shorter-lived sequences inflate their ALO, producing the observed negative correlation. Despite this reversal in sign, the magnitude of the ALO correlation still exceeds that of FE or AUP, and the strong positive correlation in the weakly paired regions (r = 0.78) provided a clear link between local structural features and mRNA stability. Overall, these results demonstrate that the LO transformation reveals structurally meaningful variation among sequences that appear similar to traditional metrics.

### Multi-feature Regression with ALO Improves mRNA Stability Prediction

Previous work by Wayment-Steele et al.^21^ evaluated statistical and deep learning models on these full-length transcripts (**Figure 1A**), achieving Spearman correlations of 0.36–0.48 on orthogonal degradation datasets^9,21^. Despite the advantage of large-scale, high-resolution input, predictive performance on full-length sequences remained modest, highlighting the challenge of generalizing across design contexts.

To test whether LO complements existing structural metrics, we incorporated ALO alongside FE/length, AUP, and GC content in a unified UMAP embedding (**Figure 5A**). This integrated model revealed improved separation between NLuc sequence clusters, with all clusters achieving statistically significant differences in in-solution half-life (**Figure 5B**). Notably, the p-value between the high half-life (red) and medium half-life (green) clusters improved from 0.62 without ALO (**Figure 1G**) to 0.023 with ALO (**Figure 5B**), based on a Mann-Whitney test representing approximately a 27-fold increase in significance. This demonstrates that LO contributes orthogonal information, enabling better resolution of stability differences that traditional global metrics alone fail to capture.

**Figure 5.**
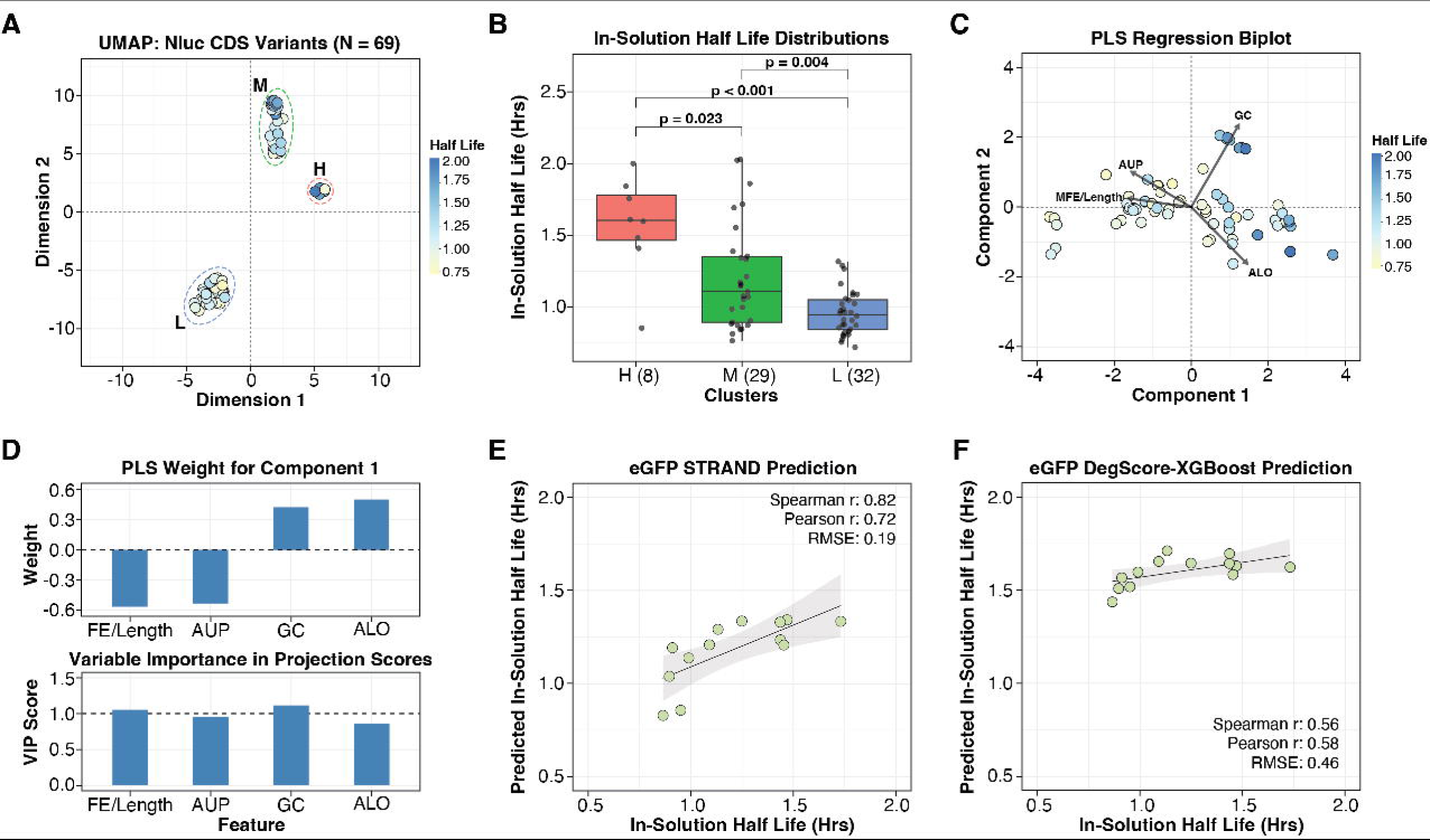
Multi-feature regression model of RNA degradation shows improved prediction accuracy through orthogonal contribution from Average Log Odds (ALO). **(A)** UMAP projection of NLuc sequences based on a multi-feature model including FE/length, AUP, GC content, and ALO, showing separation into three clusters. **(B)** Boxplot distributions of in-solution half-lives for each UMAP-derived cluster, with statistical comparisons indicating significant differences between groups. The Mann-Whitney test was used to calculate p-values. The number of sequences within each cluster is noted in parentheses. **(C)** PLSR biplot of NLuc sequences using FE/length, AUP, GC content, and ALO as features, plotted along Component 1 and Component 2. Arrows indicate loading vectors for each feature. Points are colored by in-solution half-life. **(D)** PLSR Component 1 weight of each feature (Top). Positive and negative weights indicate the direction and magnitude of each feature’s contribution to the first latent component. Variable Importance in Projection (VIP) scores across all components (Bottom). The dashed line indicates a canonically used threshold to define important variables. **(E)** PLSR model predictions of in-solution half-life for eGFP sequences using models trained on NLuc variants. A linear regression fit is shown with a 95% confidence interval represented by the shaded region. **(F)** DegScore-XGBoost model predictions for the same eGFP sequences as in **(E)**. A linear regression fit is shown with a 95% confidence interval represented by the shaded region.

Since no single metric fully explains RNA stability, we next sought to construct a multi-feature regression model integrating key structural features. An ideal model should balance explanatory power with generalizability while minimizing overfitting, particularly given the small dataset size (N = 69). Given the strong correlations among features, we selected a minimal, non-redundant feature set that captures complementary aspects of RNA structure. FE (normalized by length) and AUP represent the global secondary structure stability, which influences degradation. GC content captures sequence composition effects on stability, as GC-rich regions tend to form stronger base-pairs. ALO reflects local structural differences missed by traditional metrics, providing finer resolution of base-pairing variability. This feature set ensures that we account for thermodynamic stability, sequence composition biases, and fine-scale structural variability.

We next used Partial Least Squares Regression (PLSR)^30^ to assess the linear relationships between the feature sets and in-solution half-lives of NLuc sequences, as it is well-suited for modeling datasets with few observations and collinear predictors. The PLSR biplot revealed partially overlapping contributions from featured traditional metrics—FE normalized by length, AUP, and GC content (**Figure 5C**). GC content emerged as the dominant contributor, projecting strongly along both principal components and aligning with samples exhibiting longer half-lives. In contrast, FE/length and AUP were collinear and pointed in the opposite direction of GC, suggesting potential redundancy in the information they provide. The ALO loading vector was clearly separated from traditional metrics (both GC and FE/length), confirming its orthogonal contribution.

To further interpret the contributions of each feature in the PLSR model, we examined both the component weights and Variable Importance in Projection (VIP) scores. Component 1 weights revealed that GC content and ALO contributed most strongly to the positive direction of the latent axis associated with increasing half-life, while FE/Length and AUP contributed in the opposite direction (**Figure 5D, top**). VIP scores, which reflect the overall importance of each feature across all components, identified GC content and FE/Length as the most influential predictors (VIP = 1.05 and 1.11, respectively), with AUP and ALO contributing moderately (VIP = 0.96 and 0.86, respectively) (**Figure 5D, bottom**).

The bivariate correlations between the variables further support the unique informational content provided by ALO (**Figure S4**). While ALO exhibited strong negative correlations with FE/Length (r = −0.87) and AUP (r = −0.90), it showed moderate positive correlation with GC (r = 0.37) accompanied by a broad distribution of values. In contrast, FE/Length and AUP both showed stronger, more consistent negative correlations with GC (r = –0.66 and –0.47, respectively). This divergent relationship between ALO and GC demonstrated a new pattern of covariance that was not captured by the initial set of predictors (GC, AUP and FE/length). We refer to this sequence and structure feature-based model as Stability Regression Analysis using Nucleotide-Derived features (STRAND). STRAND integrates global (FE, AUP), local structural (ALO) and sequence features (GC) into a compact, interpretable framework for predicting mRNA degradation. We next aimed to evaluate the generalizability and predictive utility of STRAND.

Given the limited size of the NLuc dataset (N = 69), we did not perform a traditional train-test split, as doing so would risk underpowering the model and obscuring true structure-function relationships. Instead, we trained the models on the full NLuc dataset and tested their ability to generalize to the eGFP variants (N = 13). To maximize the data utilized in each training fold given our small dataset, we performed Leave-One-Out Cross-Validation (LOOCV) for model tuning and evaluation (**Figure S5A**). STRAND achieved strong predictive performance across all metrics (Spearman *r* = 0.82, Pearson *r* = 0.72, RMSE = 0.19, MAE = 0.17), highlighting the effectiveness of this compact feature set for mRNA stability prediction (**Figure 5E**).

To compare STRAND’s performance with state-of-the-art published approaches, we evaluated half-life predictions using the benchmark DegScore-XGBoost model described by Wayment-Steele et al.^21^, along with the two top-performing deep learning models from that study: Nullrecurrent, the overall first-place model, and Kazuki2, the second-place model which achieved the best performance on the specific dataset we used. For the same eGFP test set, all three models predicted degradation scores at the nucleotide level, and the average per-position degradation scores (AUP) showed strong correlation with experimental degradation rates (**Figure S5B**). Predicted half-lives for the eGFP sequences for each model were then calculated using the relation half-life = ln(2)/AUP and compared to the experimentally measured values.

The three models achieved comparable performances for the eGFP sequences, with the DegScore-XGBoost model achieving a Spearman correlation of 0.56 (Pearson r = 0.58) and RMSE of 0.46 (**Figure 5F**). NullRecurrent and Kazuki2 showed similar performance, with a Spearman correlation of 0.58 and 0.60, respectively (Pearson r = 0.40 and 0.62), and RMSE of 0.42 and 0.48 (**Figure S5C**). By comparison, STRAND achieved a Spearman correlation of 0.82, a Pearson r of 0.72, and RMSE of 0.19, outperforming all three ML models with approximately a two-fold reduction in prediction error relative to the best-performing comparison model. These results demonstrate that integrating local and global structural features in a compact model like STRAND improves both the accuracy and generalizability of mRNA degradation prediction.

To further assess STRAND’s generalizability, we applied the model to sequences derived from an independent study, VZV gE and SARS-CoV-2 Spike mRNA variants^19^. These differed substantially from the NLuc training set in both sequence length and structural feature ranges (**Figure 6A**). Moreover, in-solution half-lives of VZV and Spike sequences, measured under both 20 mM Mg^2+^ (**Figure 6A)** and 10 mM Mg^2+^ (**Figure S6A**) buffer conditions, were substantially longer than those of the NLuc training set. The LO distributions and per-position LO profiles of both VZV and Spike sequences showed a clear trend: sequences with more positions at strongly positive LO values tended to have longer half-lives (**Figures 6B, C**), consistent with the relationship between local structural confidence and stability observed in the NLuc dataset. When STRAND predictions were compared to experimentally measured half-lives, the model preserved strong rank-order accuracy for both VZV (Spearman r = 0.86) and Spike (Spearman r = 0.95) sequences, although absolute prediction errors were large (RMSE = 20.36 and 5.25, respectively) (**Figure 6D**). Notably, the FE values for both VZV and Spike sequences fall well outside the range observed in the NLuc training set, and their GC content values lie near the extremes of the NLuc distribution (**Figure 6A**). The large RMSE likely reflects extrapolation beyond the training domain and the substantial difference in half-life scale between these transcripts and the NLuc training data. This prediction trend was also maintained for the half-lives measured in 10 mM Mg^2+^ buffer condition (**Figure S6B**). Overall, these results show that STRAND generalizes across proteins in predicting stability rank order, although precise half-life prediction will require training data within the relevant distribution.

**Figure 6.**
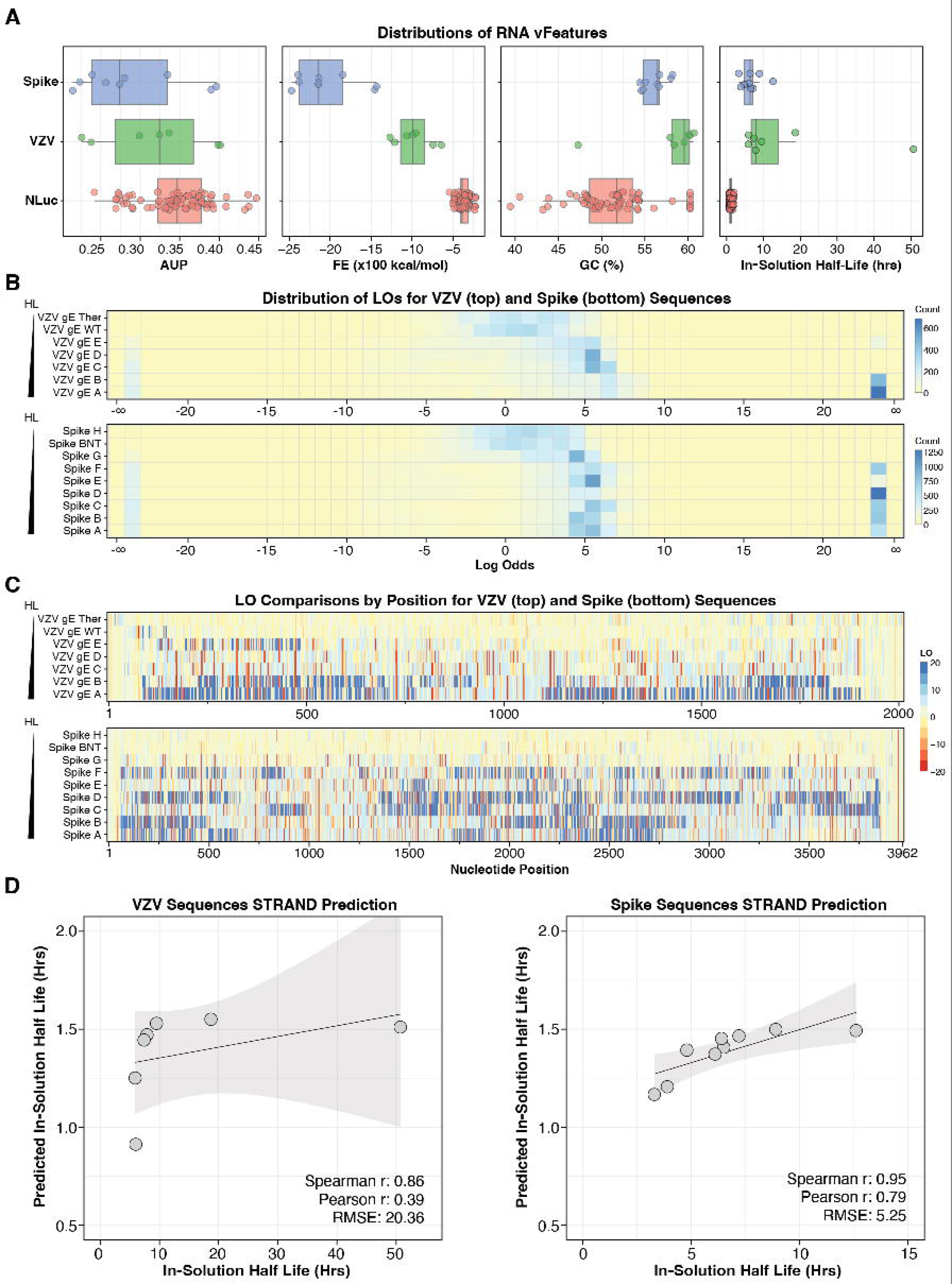
STRAND generalizes to mRNAs encoding viral antigens with structural features outside the training range. **(A)** Distributions of features (AUP, FE, GC content) and in-solution half-lives for Spike, VZV, and NLuc sequence sets, illustrating the substantial differences in feature ranges between the training (NLuc) and test (VZV, Spike) datasets. **(B)** Heatmaps of binned LO distributions (binwidth = 1) for VZV gE (top) and Spike (bottom) sequences, sorted by increasing half-life. Color scale indicates frequency of positions in each bin. **(C)** Per-position LO heatmaps for VZV gE (top) and Spike (bottom) sequences. Color scale indicates LO values. **(D)** STRAND predicted half-life versus experimentally measured half-life for VZV (left) and Spike (right) sequences. Spearman r, Pearson r, and RMSE are shown.

## DISCUSSION

In this study, we examined the relationship between RNA secondary structure and mRNA in-solution half-life, focusing on the predictive power of different structural metrics. Our findings highlight two key contributions. First, we show that ALO of base-pairing probability provides orthogonal information that complements traditional metrics such as FE, AUP, and GC content. This orthogonality enables ALO to differentiate sequences that have similar traditional structure feature values but exhibit markedly different stability. Second, we demonstrate that a multi-feature model integrating ALO with FE, AUP, and GC content outperforms previously published statistical and deep learning models in predicting mRNA stability across variants encoding multiple proteins, despite relying on a minimal and interpretable four-feature set. This compact feature combination balances complementary contributions while avoiding redundancy, resulting in a robust and generalizable framework for modeling RNA degradation.

While MFE has long been used as a proxy for RNA structural stability^12,31,32^, it provides only a single, optimal conformation^13,33^ and does not account for the diversity of structures an RNA sequence can adopt^10,11^. As illustrated in Figure 2, sequences with similar MFE values can exhibit vastly different structural topologies, such as the number and arrangement of stem-loops, bulges, or multibranch loops. These variations lead to distinct base-pairing patterns that can dramatically affect degradation rates^34,35^, yet are not captured by MFE alone.

ALO and AUP are both derived from BPP, which ranges from 0 (unpaired) to 1 (paired). While BPP provides valuable insights, it is limited by its compressed range, making it difficult to detect subtle but functionally meaningful differences near the extremes. Moreover, when BPP values are averaged across a sequence, positions with BPP near zero, which represent confidently unpaired and degradation-prone sites, are effectively treated as an absence of pairing signal rather than as informative structural features. ALO addresses this by expanding BPP values into a logarithmic scale, ranging from -∞ to +∞, thereby amplifying differences near 0 and 1 and improving interpretability of pairing confidence^26^. For example, BPP values of 0.9, 0.99, and 0.999 are all close to 1 and difficult to distinguish on the original scale, but map to log-odds of 2.20, 4.60, and 6.91, respectively, offering clearer resolution of high pairing confidence. Similarly, low BPP values (e.g., 0.01 vs. 0.1) become more distinguishable once transformed, allowing better identification of confidently unpaired regions. This enhanced contrast allows ALO to highlight structural differences between sequences that appear similar using traditional metrics, as demonstrated by our binning analysis. These findings led us to explore whether combining ALO with other established metrics could enhance predictive modeling of mRNA stability.

Building on these insights, we developed STRAND, a multi-feature regression model incorporating ALO, FE, AUP, and GC content. Despite its compact four-feature design, this model performs competitively with previously published statistical and deep learning approaches while maintaining greater interpretability. Importantly, it generalizes well to independent data, including eGFP, VZV gE, and SARS-CoV-2 Spike constructs, with strong rank-order performance even when applied to transcripts well outside the training domain. The model’s simplicity makes it especially suitable for early-stage screening and optimization workflows, where interpretability and low feature complexity are critical. Moreover, its performance without reliance on experimental probing data highlights the value of well-chosen *in silico* features for robust prediction.

Beyond its predictive utility, STRAND and the LO metric can be readily incorporated into existing mRNA design workflows (**Figure S7**). During the initial sequence design stage, LO or ALO can be an additional objective or constraint within optimization algorithms, complementing traditional metrics to guide the sequence generation with favorable local structural profiles. Alternatively, for a given candidate sequence pool, STRAND can be used to predict in-solution half-lives, or ALO can serve as a filter to prioritize candidates with structural features associated with greater stability prior to experimental validation. In either application, STRAND’s computational simplicity and its reliance on readily computed *in silico* features make it practical for integration into high-throughput design pipelines.

While STRAND performs well overall, there are limitations that offer opportunities for further improvement. As expected, prediction accuracy declines at the extremes of the half-life range, likely due to limited representation of highly stable or unstable sequences in the training set (**Figure 1A**). One of the most influential biological variables affecting degradation is transcript length^7,36–38^, yet STRAND was trained using variants of a single open reading frame (ORF), as it was the only transcript in the dataset with a sufficient number of sequence variants for training. While STRAND performed reasonably well even for sequence sets of different length and feature values outside of the training regime, expanding the training dataset to include transcripts encoding proteins of varying lengths and sequence features would improve the model’s robustness and generalizability. For example, STRAND predictions for MEV sequences were poor (**Figure S8**). This is likely attributable to the short transcript length of MEV variants, for which FE normalization by length becomes unreliable due to the non-linear relationship between FE and sequence length at shorter lengths^39^, combined with their narrow FE distributions that fall outside the training regime. Increasing the diversity of the dataset may uncover new features that contribute independently of both ALO and MFE, providing further gains in predictive accuracy and biological insight. Moreover, STRAND currently models CDS-level structural features and does not explicitly account for other important dimensions of mRNA design, such as co-optimization of UTR features, or incorporating structure predictions specific for relevant modified nucleotides. Investigating how these factors interact with the structural features captured by STRAND represents an important direction for future work.

For any given value of conventional metrics such as MFE, sequences often share similar range of LO values but differ in their distributions, making it difficult for LO alone to distinguish stability differences across broader structural contexts. LO effectively resolves differences between sequences that share similar global metrics but differ in degradation rates, capturing subtle structural features that traditional approaches often overlook. However, it lacks the broad discriminatory power needed to classify sequences across a wider spectrum of structural contexts. Therefore, the strength of LO lies in complementing existing metrics, providing finer-scale insights particularly valuable in cases where traditional metrics offer limited differentiation.

Additionally, a key limitation of our approach stems from inherent inaccuracies in *in silico* RNA secondary structure predictions, which rely on imperfect thermodynamic parameters and do not always fully capture biologically relevant structural ensembles^40–42^. As our stability model relies on these predicted structures, its accuracy is likely influenced by the quality of the folding algorithms. Continued improvements in RNA structure prediction methods could therefore further enhance the performance and reliability of STRAND.

Across the NLuc, eGFP, VZV, and Spike datasets, we observed distinct patterns of LO distributions that are associated with differences in degradation. Understanding how the balance between highly structured and weakly structured regions contributes to degradation across different sequence contexts remains an open question. Nevertheless, STRAND demonstrates that well-chosen structural features can substantially reduce prediction error for mRNA stability, and its compact, interpretable design makes it readily adaptable as new training data and design contexts become available.

## METHODS

### RNA Structure Prediction and Metric Calculation

#### Data Acquisition

RNA sequence and corresponding half-life data were obtained from the study by Leppek et al.^9^ accompanying the OpenVaccine: COVID-19 mRNA Vaccine Degradation Prediction dataset^9,21^.

### Feature Calculation using LinearPartition

Acquired RNA sequences were used as input for the LinearPartition^29^ software to generate structural metrics, including the FE predictions, the BPP for each nucleotide position, the average base pairing probability across the sequence, and the AUP across the sequence. The BPP at each nucleotide position is calculated as the summed probability of that nucleotide forming a base pair with any other nucleotide across the entire predicted structural ensemble derived from the partition function.

### Structure Visualization using RNAfold

For visual representation of predicted RNA secondary structures, sequences were submitted to the RNAfold^43^ web server. Default options provided by the web server were used for both structure prediction and visualization.

### Log Odds Calculation

Using the BPP calculated by LinearPartition^29^, the Log Odds Per Position (LOPP) was determined for each nucleotide using the formula: LOPP = ln(p / (1-p)), where p is the BPP. To handle probabilities of exactly 0 or 1, a pseudocount of 1e^-10^ was added to p when BPP = 0 and subtracted from p when BPP = 1 prior to the log odds calculation.

### BPP Curvature Analysis and Thresholding

The relationship between BPP and LO was examined by calculating the mathematical curvature of the BPP vs. LO curve. Curvature was determined by numerically calculating the second derivative of LO with respect to BPP (d²(ln(p/(1-p)))/dp²) within the boundary conditions 0 ≤ p ≤ 1. This analysis identified points of maximum curvature at BPP values of approximately 0.20 and 0.80. Based on this, a BPP threshold of 0.2 was selected to define positions considered likely to be unpaired for downstream analyses.

### Sequence Clustering Analysis

Uniform Manifold Approximation and Projection (UMAP)^28^ was employed to explore potential clustering based on sequence features. Input features included FE normalized by sequence length, Average Unpaired Probability (AUP), and GC content. Analyses were performed both including and excluding the Average Log Odds (ALO) as an additional feature. UMAP was performed using the UMAP package (Konopka, T. ; DOI: <u>10.32614/CRAN.package.umap</u>) in R (v 4.4.0 ; https://www.r-project.org) utilizing default parameters (including n_neighbors=15, min_dist=0.1, n_epochs=200). Due to the stochastic nature of the algorithm, UMAP analysis was repeated 20 times for each feature set; cluster consistency was assessed by visually confirming the presence of three distinct clusters and the stability of sequence identities within each cluster across runs.

### Linear Regression Modeling

Linear regression models using FE and AUP as predictors were generated using the stats package in R (v 4.4.0).

### PLS Regression Modeling

To model the relationship between sequence-derived features and in-solution half-life, Partial Least Squares Regression (PLSR)^30,44^ was implemented using the pls package in R (v 4.4.0). The final set of predictor variables included FE normalized by sequence length, Average Unpaired Probability (AUP), GC content, and Average Log Odds (ALO). Predictor variables were centered and scaled prior to modeling. Given the limited training dataset size (N = 69), Leave-One-Out Cross-Validation (LOOCV) was utilized on the training set for model parameter tuning by minimizing the Root Mean Squared Error of Prediction (RMSEP) (Fig. S5A). Variable Importance in Projection (VIP) scores were calculated using the plsVarSel package^45^ in R (v 4.4.0 ; https://www.r-project.org) to assess the contribution of each predictor variable to the final PLSR model trained on the full training set. Final model performance was evaluated on a separate, independent EGFP dataset (N = 13) by calculating the Pearson and Spearman correlation coefficients, and Root Mean Square Error (RMSE) between predicted and actual half-life values.

## DATA AVAILABILITY

The STRAND model described in this paper is publicly available to support reproducibility and broad use. The raw training data used in this study were obtained from the OpenVaccine: COVID-19 mRNA Vaccine Degradation Prediction dataset (Leppek et al., 2022). Any additional data required to verify and extend the research are available from the corresponding author upon request.

## AUTHOR CONTRIBUTIONS

**S.Y.:** Conceptualization (Lead), Formal analysis (Lead), Investigation (Lead), Validation (Lead), Visualization (Lead), Writing – original draft (Lead), Writing – review & editing (Lead). **S.E.A.:** Data curation (Supporting), Formal analysis (Supporting), Investigation (Supporting), Writing – original draft (Supporting), Writing – review & editing (Supporting). **Y.J.:** Formal analysis (Supporting), Investigation (Supporting), Validation (Supporting), Writing – review & editing (Supporting). **J.W.D.:** Conceptualization (Lead), Supervision (Supporting), Writing – review & editing (Supporting). **M.M.:** Conceptualization (Lead), Investigation (Lead), Project administration (Lead), Resources (Lead), Supervision (Lead), Writing – original draft (Lead), Writing – review & editing (Lead).

## ACKNOWLEDGEMENTS

The authors thank Chris Pepin for the preliminary discussions of this work, Alicia Bicknell and Chiaowen Joyce Hsiao for critical reading of the manuscript. This work was supported in part by the NIH under the award T32 GM152319 (S.Y.)

## Declaration of generative AI and AI-assisted technologies in the manuscript preparation process

During the preparation of this work the author(s) used Claude and ChatGPT to assist with drafting and editing portions of the manuscript. After using this tool/service, the author(s) reviewed and edited the content as needed and take(s) full responsibility for the content of the published article.

