## Supplemental Figures for "Fine-Scale Structural Information Substantially Improves mRNA Therapeutic Stability Prediction"

### Figure S1

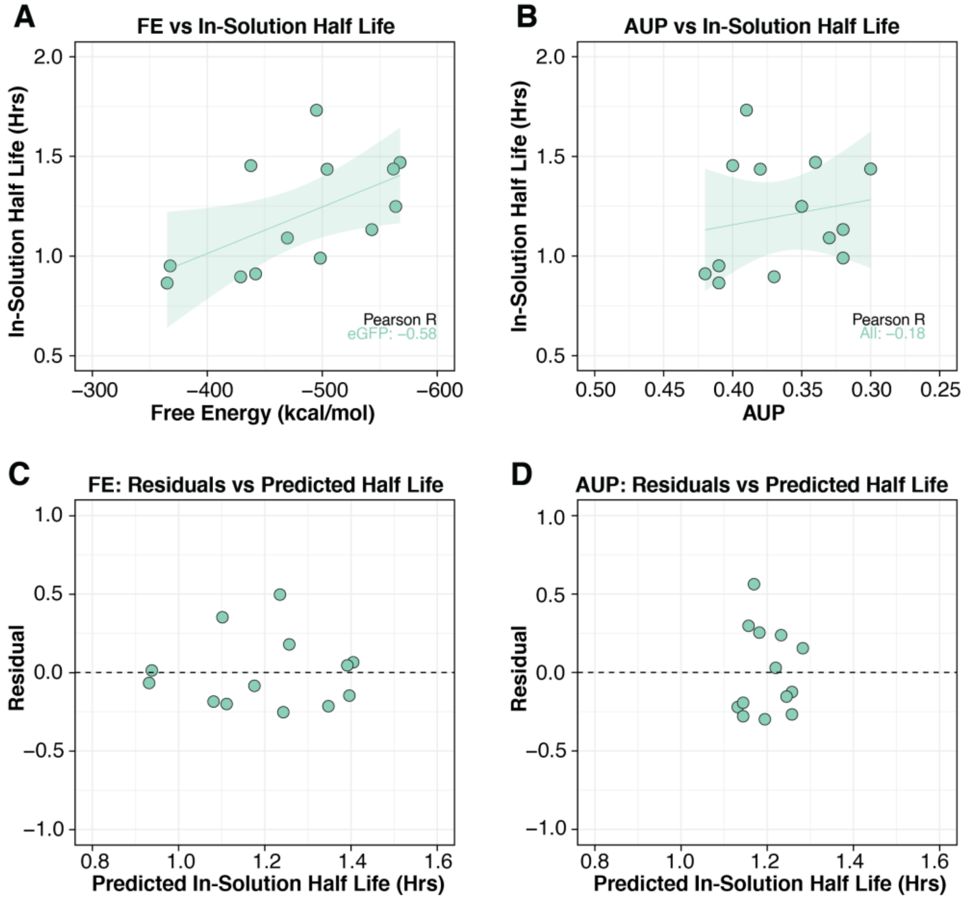


**Figure S1. Structure–function analysis in the eGFP set. (A)** Correlation of Free Energy (FE) and half-life in the eGFP dataset. **(B)** Same as (A) but for AUP. **(C)** Residual plots for FE vs in-solution half-life. **(D)** Same (C) but for AUP.

**Figure S2**

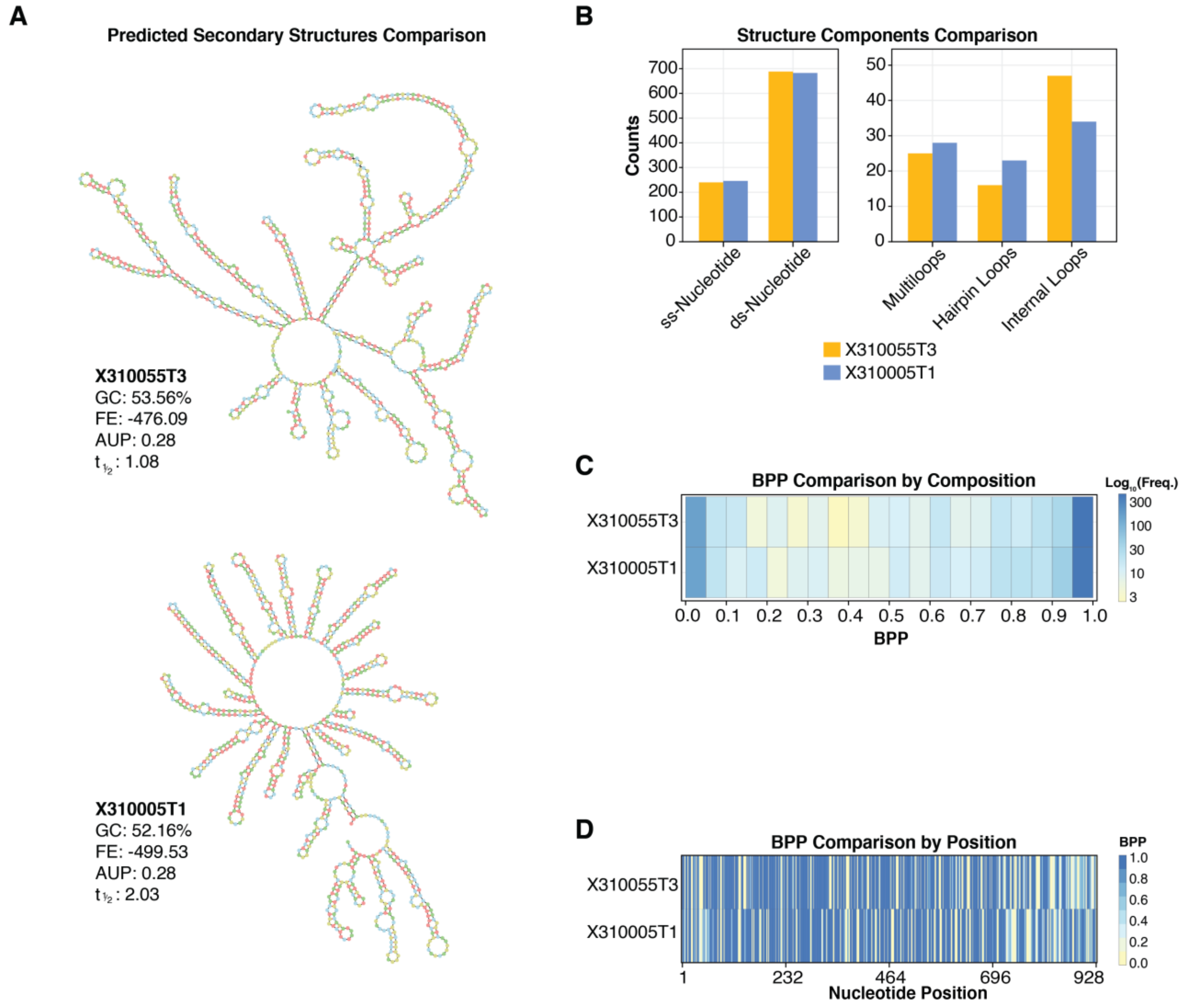


**Figure S2. Structure comparison between two RNA sequences at lower FE bin with similar global metrics but different in in-solution half-life (A)** Predicted secondary structures of two NLuc sequences, X310055T3 and X310005T1, which exhibit similar global structure metrics (FE and AUP) but differ substantially in in-solution half-life. **(B)** Barplot comparing structural components between the predicted maximum expected accuracy structures, including counts of unpaired nucleotides, hairpin loops, interior loops, and multiloops. **(C)** Heatmap comparing BPPs binned at 0.05 resolution, showing clear differences in the distribution of local pairing strength between the two sequences. **(D)** Heatmap comparing local BPP values by position, revealing fine-scale structural differences between the two sequences.

**Figure S3**

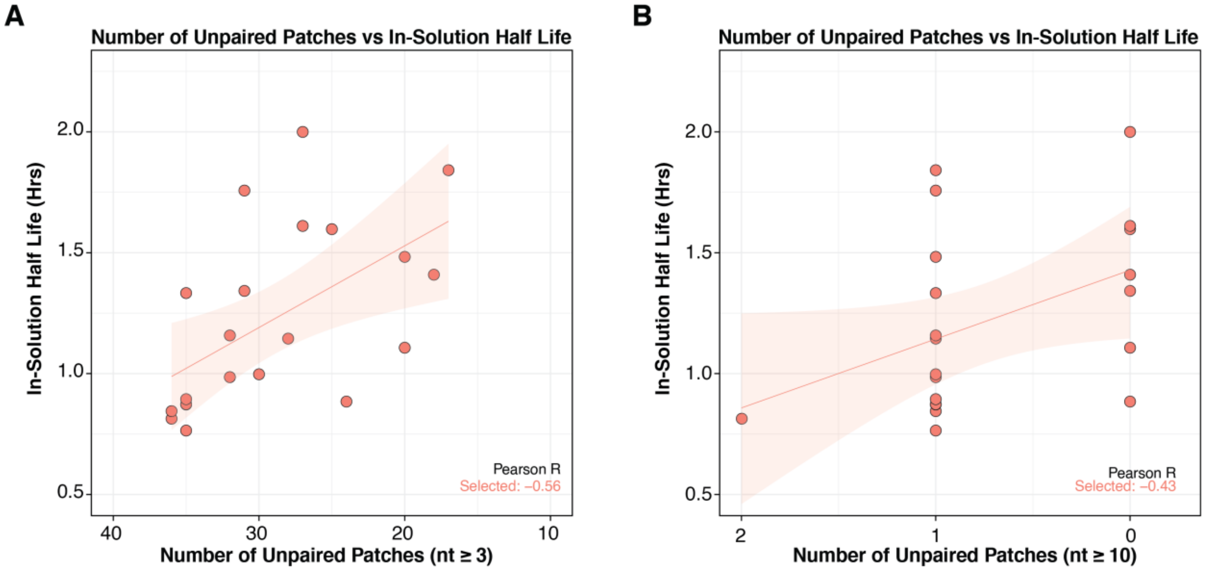


**Figure S3. Correlation between the number of unpaired patches and in-solution half-life (A)** ≥3 consecutive unpaired positions with BPP < 0.2 **(B)** ≥10 consecutive unpaired positions with BPP < 0.2.

**Figure S4**

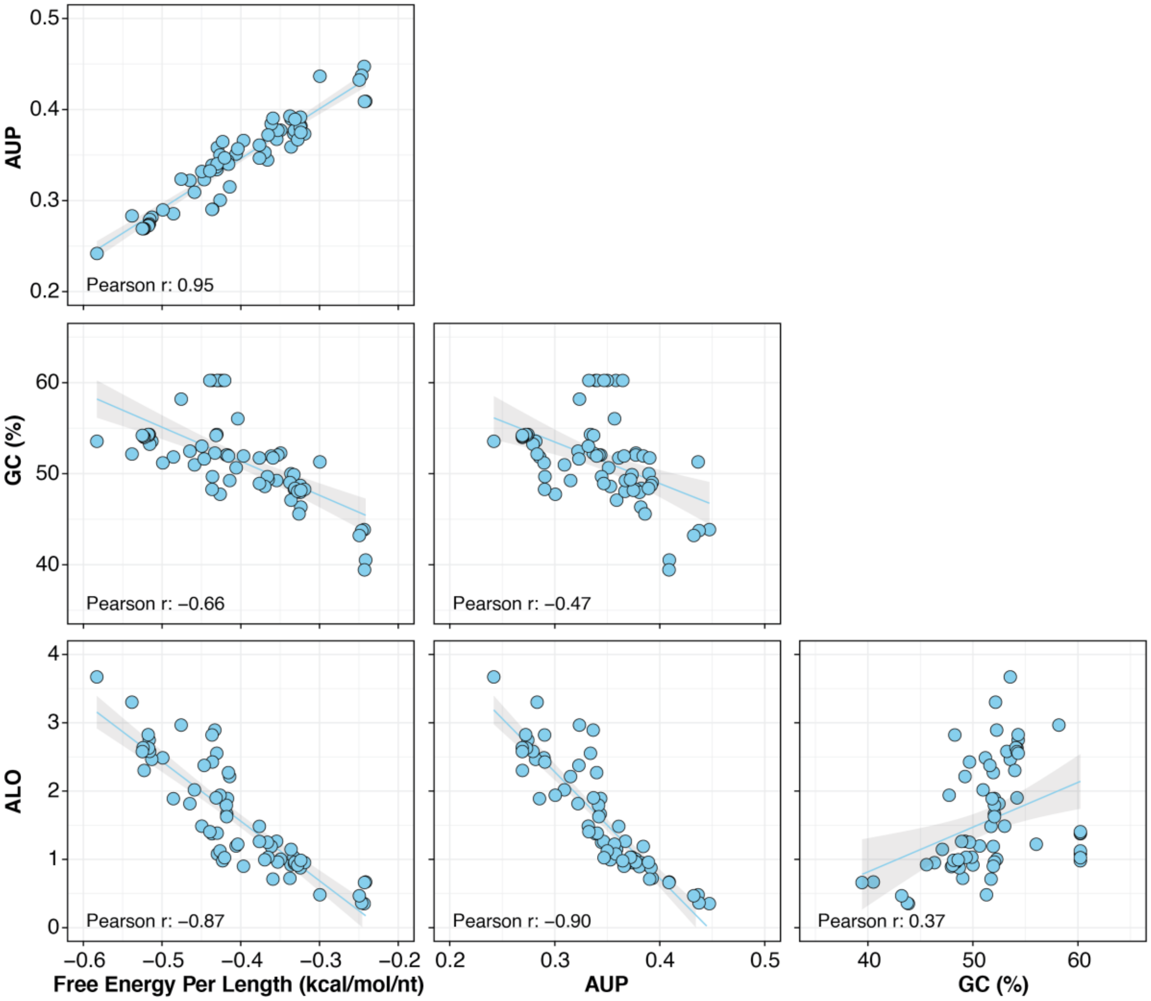


**Figure S4. Bivariate correlations between independent variables used in PLSR modeling.** The relationships between the independent variables (FE/Length, AUP, GC, and ALO) for the curated NLuc mRNA variants[1] used in PLSR modeling are shown.

**Figure S5**

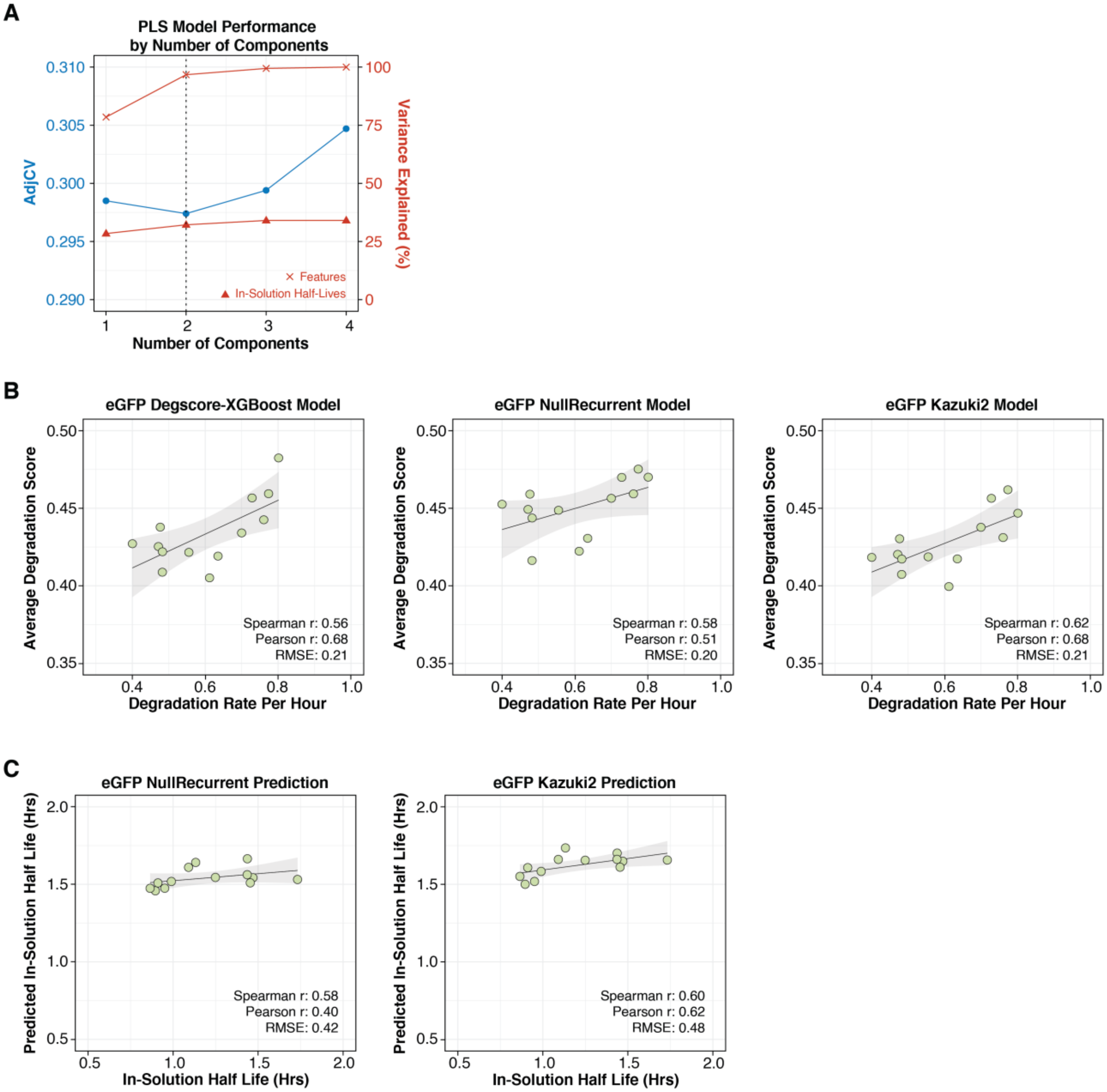


**Figure S5. PLS Model Parameters and Comparison Model Predictions. (A)** PLS model performance by number of components, showing AdjCV (blue) and variance explained (red) for features and in-solution half-lives. The dashed line indicates the selected number of components. **(B)** Average per-position degradation scores versus experimental degradation rate per hour for eGFP sequences, shown for DegScore-XGBoost, NullRecurrent, and Kazuki2 models. Spearman r, Pearson r, and RMSE are shown for each. **(C)** Predicted in-solution half-life (calculated as ln(2)/AUP) versus experimentally measured half-life for NullRecurrent and Kazuki2 models.

**Figure S6**

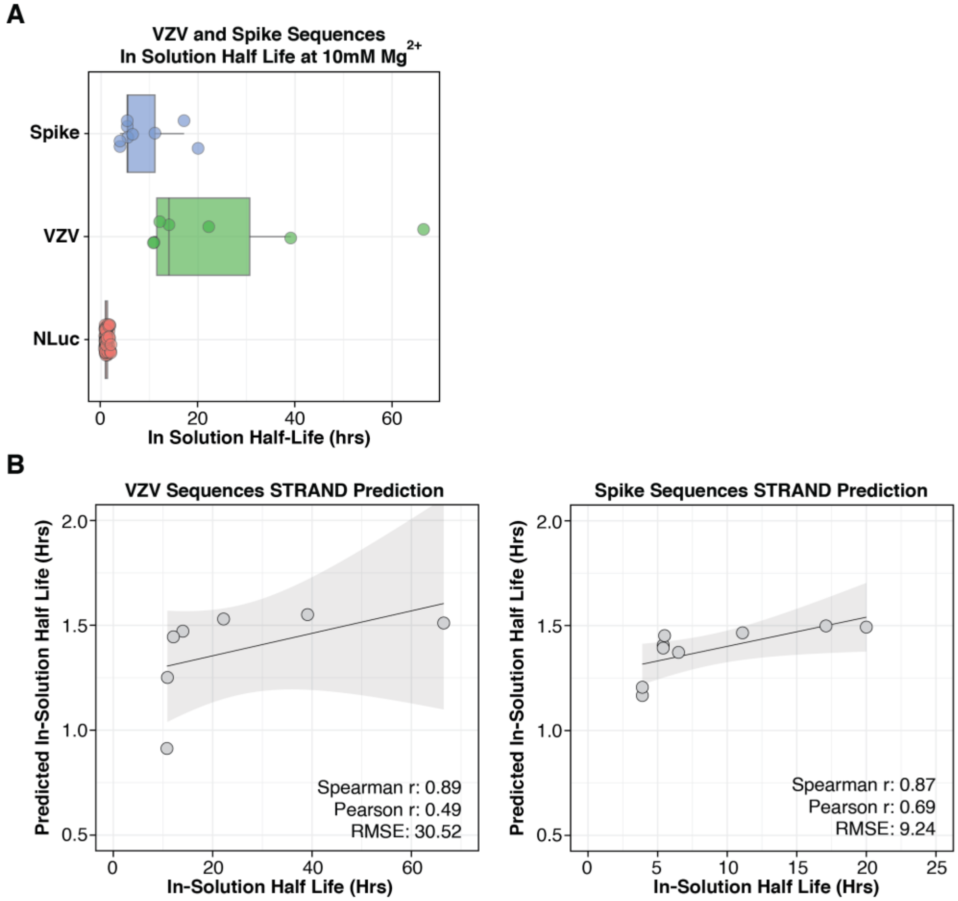


**Figure S6. Application of STRAND to VZV and Spike Sequences with Measured In-Solution Half Lives at 10 mM $Mg^{2+}$ buffer conditions. (A)** Distribution of in-solution half-lives for Spike and VZV sequences measured at 10 mM $Mg^{2}$+. Half-lives of NLuc training set are also shown. **(B)** STRAND predicted half-life versus experimentally measured half-life for VZV (left) and Spike (right) sequences. Spearman r, Pearson r, and RMSE are shown.

**Figure S7**

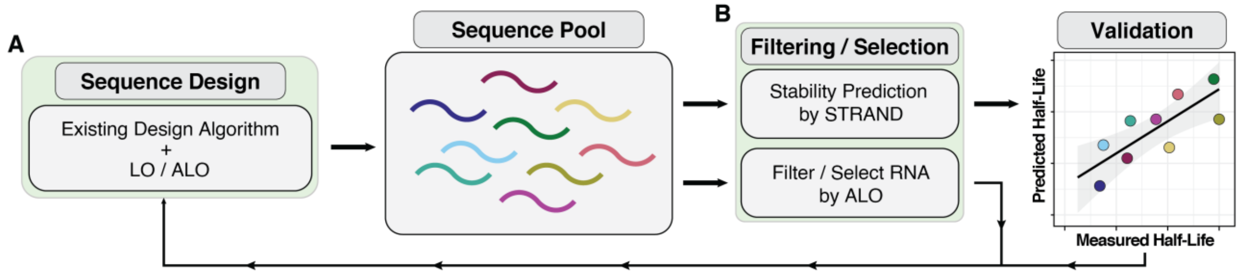


**Figure S7. Framework for applying ALO and STRAND in rational mRNA design.** Schematic describing how ALO and STRAND can be incorporated into mRNA design workflows. **(A)** LO or ALO can be integrated as an additional objective or constraint within existing mRNA design algorithms during sequence generation. **(B)** For a given pool of candidate sequences, STRAND can be used to predict in-solution half-lives or ALO can be used to filter and prioritize candidates with favorable local structural profiles prior to experimental validation.

**Figure S8**

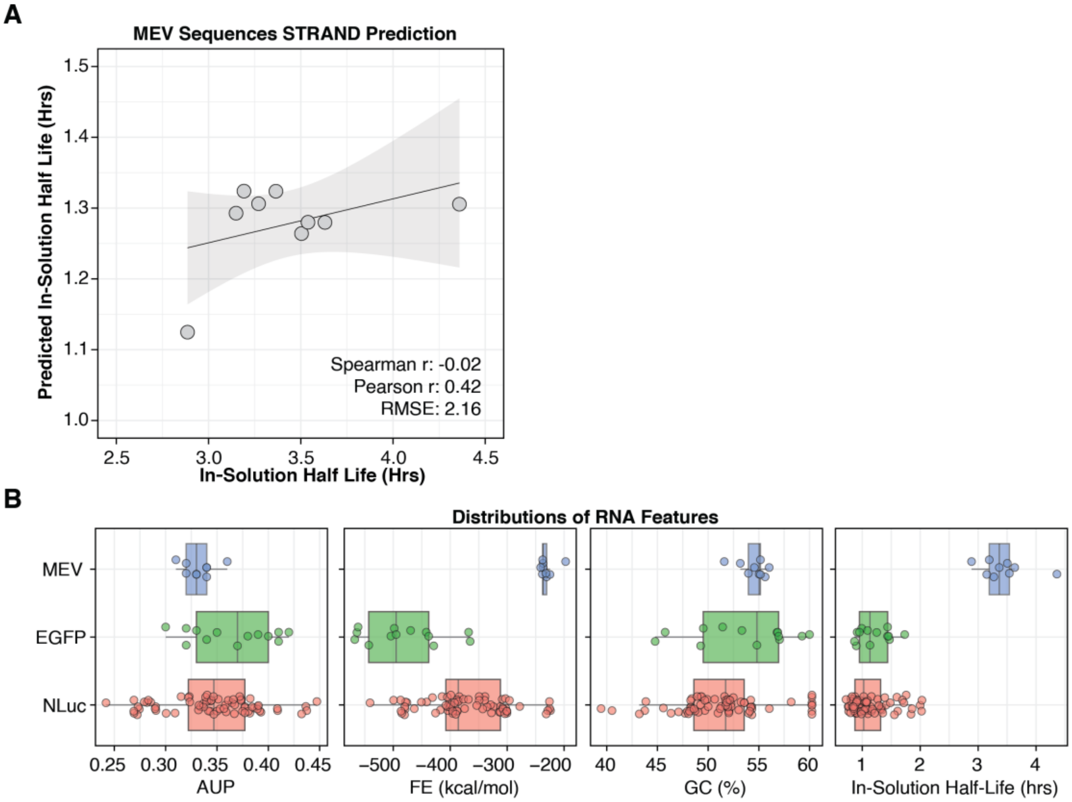


**Figure S8. STRAND Applied to MEV Sequences. (A)** STRAND predicted half-life versus experimentally measured half-life for MEV sequences. Spearman r, Pearson r, and RMSE are shown. **(B)** Distributions of structural features (AUP, FE, GC content) and in-solution half-lives for MEV, eGFP, and NLuc sequence sets.
